# Assessing the influence of distinct IVF culture media on human pre-implantation development using single-embryo transcriptomics

**DOI:** 10.1101/2022.10.05.510961

**Authors:** Bastien Ducreux, Julie Barberet, Magali Guilleman, Raquel Pérez-Palacios, Aurélie Teissandier, Déborah Bourc’his, Patricia Fauque

## Abstract

The use of Assisted Reproductive Technologies (ART) is consistently rising across the world. However, making an informed choice on which embryo culture medium should be preferred to ensure satisfactory pregnancy rates and the health of future children critically lacks scientific background. Particularly, embryos within their first days of development are highly sensitive to their micro-environment. Here, we aimed to determine the impact of culture media composition on gene expression of human preimplantation embryos. By employing single-embryo RNA-sequencing after 2 or 5 days of post-fertilization culture in different commercially available media, we revealed medium-specific differences in gene expression changes. In particular, we found that culture medium composition can affect the dynamics of expression of developmentally relevant genes at day-2 but the differences were mitigated at the blastocyst stage. This study highlights the ability of embryos conceived in suboptimal *in vitro* culture media to recover proper transcriptome competency.

## Introduction

Assisted reproductive technologies (ARTs) have allowed the birth of millions of children worldwide. In Europe for instance, above two millions of children were born following ART ^1^ and numbers continuously rise, proving that tackling infertility is a huge challenge for decades to come ^2^. However, significant variability in ART practice and effectiveness exists between countries, and even at the regional scale ^3,4^. In particular, embryo culture is at the core of ART, but making an informed decision on which culture medium to use is still an arduous task ^5^. Embryo culture media are not expected to perfectly mirror *in vivo* environment conditions ^6^, but they should nonetheless provide the required biological content to sustain satisfactory embryo development compared to natural conceptions. A myriad of embryo culture media is nowadays commercially available. However, owing to trade confidentiality, their exact composition is unknown, which obscures the scientific decisions for choosing one culture medium over another for embryologists ^7^. Although the competitive commercial race to optimize embryo culture media greatly contributed to increase pregnancy rate in ART, the scientific basis behind their formulation is unclear, which is a matter of concern for ART-related biovigilance ^8^. Very few studies have followed up the health of children born in relation to different embryo culture media used in ART cycles, but they tend to indicate that certain media may be suboptimal, with potential long-term health impact ^9–11^.

The early embryo closely interacts with its environment, particularly during the cleavage stage ^12,13^. After fertilization, the embryo transits through the oviduct until reaching the uterus where it may implant. This journey throughout the maternal track exposes the embryo to multiple molecules, including growth factors, hormones and metabolites, which promote complex reactions ^14–16^. This period also coincides with critical epigenetic reprogramming, which influences gene expression ^17,18^. Substantial evidence have linked adverse environmental maternal exposures and transcriptome changes in human embryonic stem cells and newborns’ cord blood ^19,20^. This likely reflects the adaptation of the embryo to external stressors, displaying remarkable plasticity at the molecular and cellular level ^21,22^.

Compared to natural conception, *in vitro* conditions inherent to ART can be a source of additional stress ^23^. Indeed, the *in vitro* environment has been suspected of introducing detrimental abnormalities in the population born *in vitro* ^24–26^. The many processes involved in ART and in particular the osmotic stress, substrate imbalance, volatile organics and contaminant pollution linked to *in vitro* culture can alter fundamental embryonic biological mechanisms ^27–29^. As Thompson *et al*. ^30^ pointed out, “there is no adaptation by an embryo to its environment that has no consequence”. In particular, differences in embryo culture medium composition may lead to differences in adaptive responses to stress.

Until embryonic genome activation (EGA), the embryo is mostly transcriptionally silent and relies on maternally provided mRNAs ^31^. During this period, the capacity of the embryo to maintain metabolism and cellular homeostasis may thus be limited ^32,33^. Accordingly, short exposure to ammonium before compaction was shown to compromise the ability to further develop compared with the same exposure after compaction in mouse ^12^. After EGA, dynamic changes of gene expression accompany embryonic lineage specification, and anomalies in these sequential expression changes can lead to developmental arrest ^34^. These examples highlight that the early embryo is sensitive to its micro-environment and that many parameters in *in vitro* fertilization (IVF) centers should be tightly controlled, especially embryo culture.

Evidence from animal models showed that *in vitro* culture can affect embryonic gene expression and epigenetic marks compared with *in vivo* conditions ^35–38^. Most importantly, these molecular effects can be worsened depending on the culture medium ^36,39,40^, with a reported sensitivity of imprinted gene expression ^22,41^. Comparatively, studies in humans have mainly focused on the clinical efficiency of various culture media (live birth rate, implantation rate, clinical pregnancy rate, birthweight, placental weight, preterm birth rate) ^42–45^. Only two studies compared the transcriptomic profile of blastocysts cultured in two different media: using microarrays, they reported misregulation of genes involved in cell-cycle, apoptosis, protein degradation and metabolism, which has the capacity to impair embryo development ^48,49^. This, coupled with the observation that ART embryos have lower quality than naturally conceived embryos ^22^, justifies to pursue efforts to identify the biological origin of embryo culture effects.

In this study, we investigated for the first time the impact of different culture media used in IVF centers and known to differ in pre-implantation development performance on the transcriptome of day-2 and day-5 human embryos using single-embryo RNA-sequencing. We found evidence for medium-specific changes at day-2, affecting major genes involved in embryonic development. However, differences tended to reduce upon extended culture until blastocyst stage (day-5), reflecting the possible adaptation of the embryonic transcriptome to culture medium. Finally, establishing an expression pseudotime, we evaluated whether the dynamics of genes contributing to embryonic development could be affected by culture medium composition.

## Results

### Study design and quality control analysis by comparison with previous studies

To analyze the impact of embryo culture media on the embryonic transcriptome, we performed single embryo RNA-seq ^46^ on 51 vitrified/warmed donated embryos after day-2 or day-5 of culture (Figure 1). We compared three different media: Global (LifeGlobal, USA), SSM (Irvine Scientific, USA) and Ferticult (FertiPro, Belgium). Global and SSM have very similar compositions, except different forms of glutamine and the presence of taurine in SSM ^11^, and are intended to be used as one-step media up to day-5/6 of human embryo development. Ferticult differs from both in that it does not contain amino acids, and is fitted to be used up to day-2/3. We processed 31 day-2 embryos (SSM: *n*=14, Global: *n*=11 and Ferticult: *n*=6) and 20 day-5 embryos (Global: *n*=13 and Ferticult: *n*=7). An average of 3.3 million reads per embryo at day-2 and 13.1 million reads at day-5 was generated, with an average mapping rate of 90.1% across all samples (Supplementary Table S1). We were able to detect the expression of 30% and 26% of all RefSeq genes and transposable elements at day-2 and day-5, respectively.

**Figure 1.**
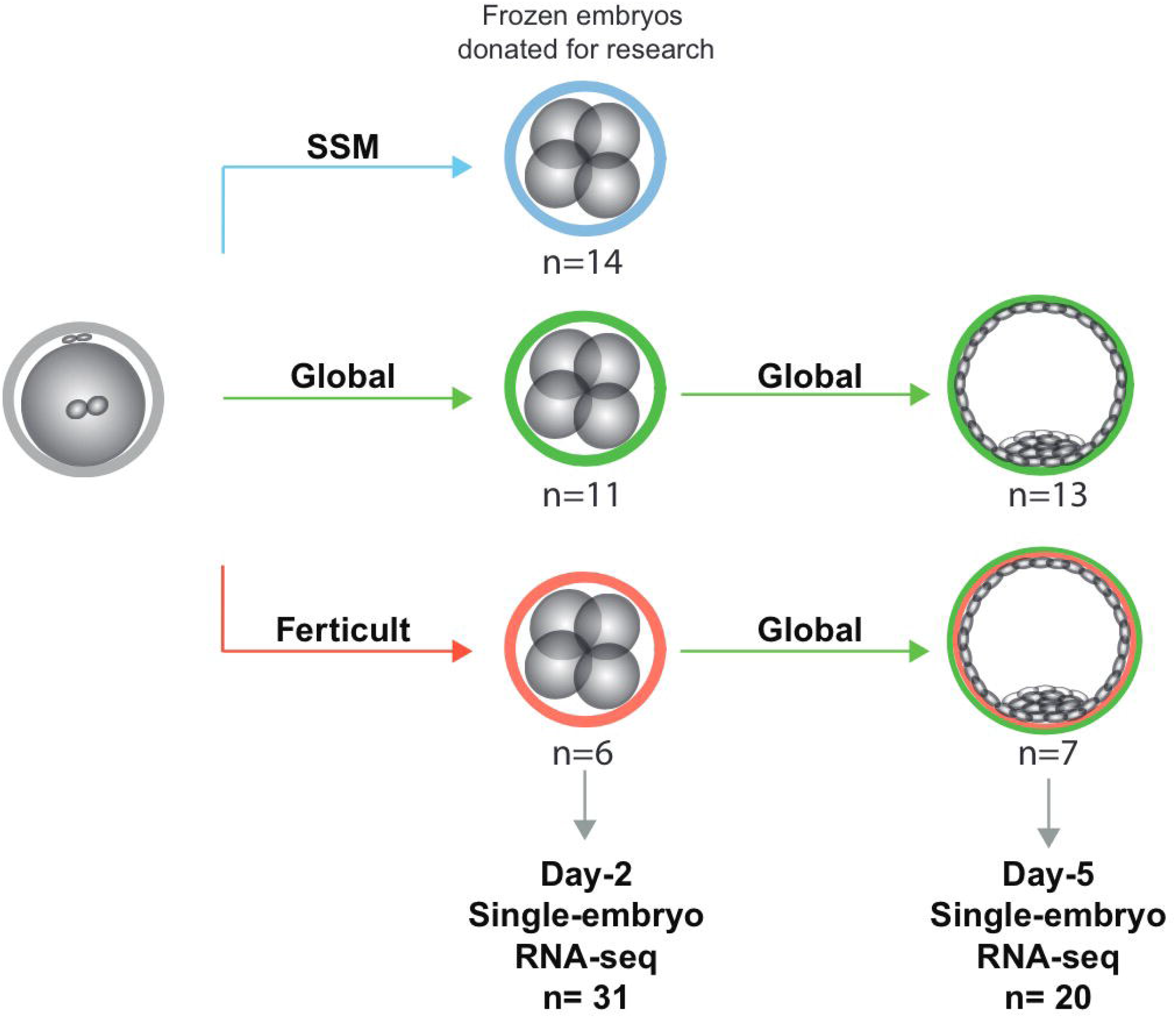
Study design for the transcriptomic analysis of human embryos cultured until day-2 and day-5. At day-5, the number of samples was independent of the rate of embryos that survived the thawing process and reaching at least the B2 stage (42.3% and 50.0% in Global and Ferticult groups, respectively).

To assess the quality of our generated single-embryo RNA-seq datasets, we relied on previous high-quality studies that performed single-cell RNA-seq (scRNA-seq) in human early embryos. According to the criteria that an expressed gene should have a count per million (cpm) value greater than 1 in at least half of the samples in each embryo stage, we found consistent numbers of 12,056, 10,022 and 11,213 genes being expressed in 4-cell stage embryos in Yan *et al*. ^47^, Xue *et al*. ^48^ and our own dataset, respectively (Supplementary Figure S1). In blastocysts, we identified 12,790 expressed genes in our data, compared with 8,204 genes in Yan *et al*. dataset ^47^ in which blastocysts were collected a day later, at day-6. Principal component analysis (PCA) of global gene expression further confirmed the high similarity of our data with those two previous studies: our day-2 embryo samples clustered near 4-cell samples and our day-5 embryos samples clustered with late blastocysts (Supplementary Figure S2).

### Transcriptomic comparison of day-2 embryos cultured in different media

We first focused on transcriptome differences on short-term culture, at the 4-cell stage (day-2), between embryos conceived in the different media. We performed differential expression for each gene expressed in at least 4 samples comparing each of the three culture media groups one by one. The highest number of differentially expressed genes (DEGs) was found comparing Ferticult and Global media groups, with 266 DEGs showing adjusted *p*-value<0.1 (Figure 2A and Supplementary Table S2). In contrast, only 1 and 5 DEGs were found comparing SSM with either Ferticult or Global groups, respectively (Supplementary Table S2). However, global transcriptomic analysis showed that SSM was transcriptionally closer to Global than to Ferticult (r=0.95 versus r=0.9, Spearman’s correlation), which is consistent with their similar composition (Supplementary Figure S3).

**Figure 2.**
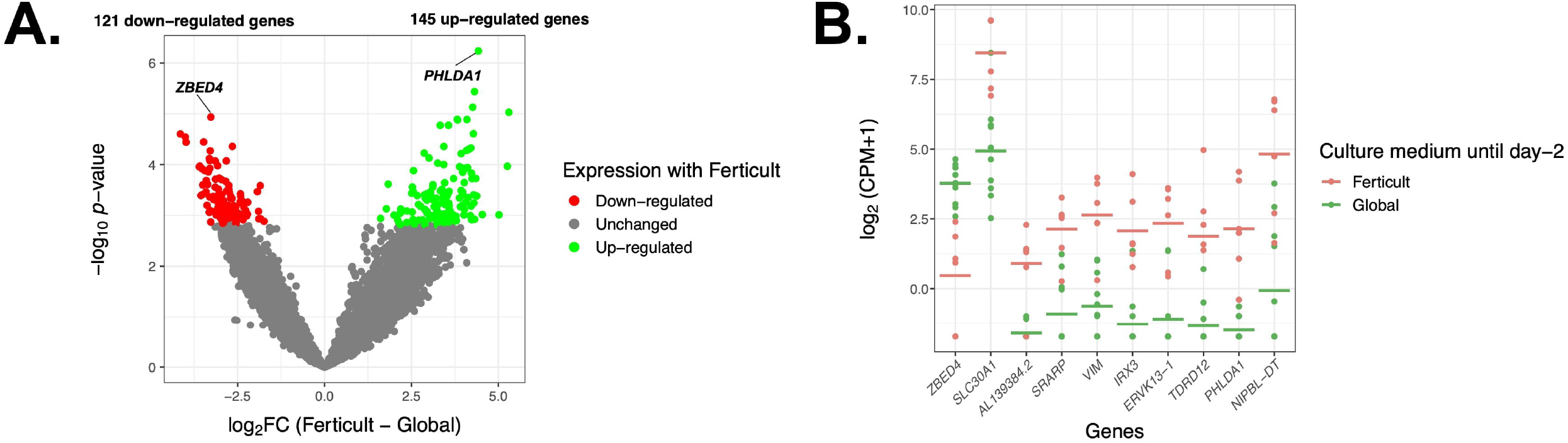
Differential gene expression analysis in day-2 embryos between Ferticult and Global media. A. Volcano plot of gene expression between Ferticult and Global media at day-2. B. Dot plot of the expression of top-10 Ferticult-to-Global DEGs, ordered by ascending log2FC (from left to right). Group mean is represented by the line. Dots represent individual embryos.

Among the 266 DEGs in the Ferticult-to-Global comparison, 145 were up-regulated and 121 down-regulated. Most of them (88.3%) displayed absolute log2(Fold-Change, FC))>2.5, which revealed substantial differences in the transcriptome of day-2 embryos depending on the culture medium (Supplementary Table S2). Top-10 most dysregulated genes included *ZBED4, SLC30A1, AL139384*.*2, SRARP, VIM, IRX3, ERVK13-1, TDRD12, PHLDA1* and *NIPBL-DT* (Figure 2B, Supplementary Table S2). Gene Ontology (GO) analysis with Metascape revealed an over-representation of DEGs related to embryonic development (pattern specification process, reproductive structure development, skeletal system development, kidney development), regulation (regulation of mitotic cell cycle, regulation of cyclin-dependent protein kinase activity, negative regulation of intrinsic apoptotic signaling pathway in response to DNA damage, positive regulation of transforming growth factor), ribonucleoprotein biogenesis complex and response to nutrient (Figure 3A, 3B and Supplementary Figure S4). In parallel, gene set enrichment analysis (GSEA) revealed the top-10 significant ontologies were all related to embryonic development and cell division (Figure 3C, Supplementary Table S3).

**Figure 3.**
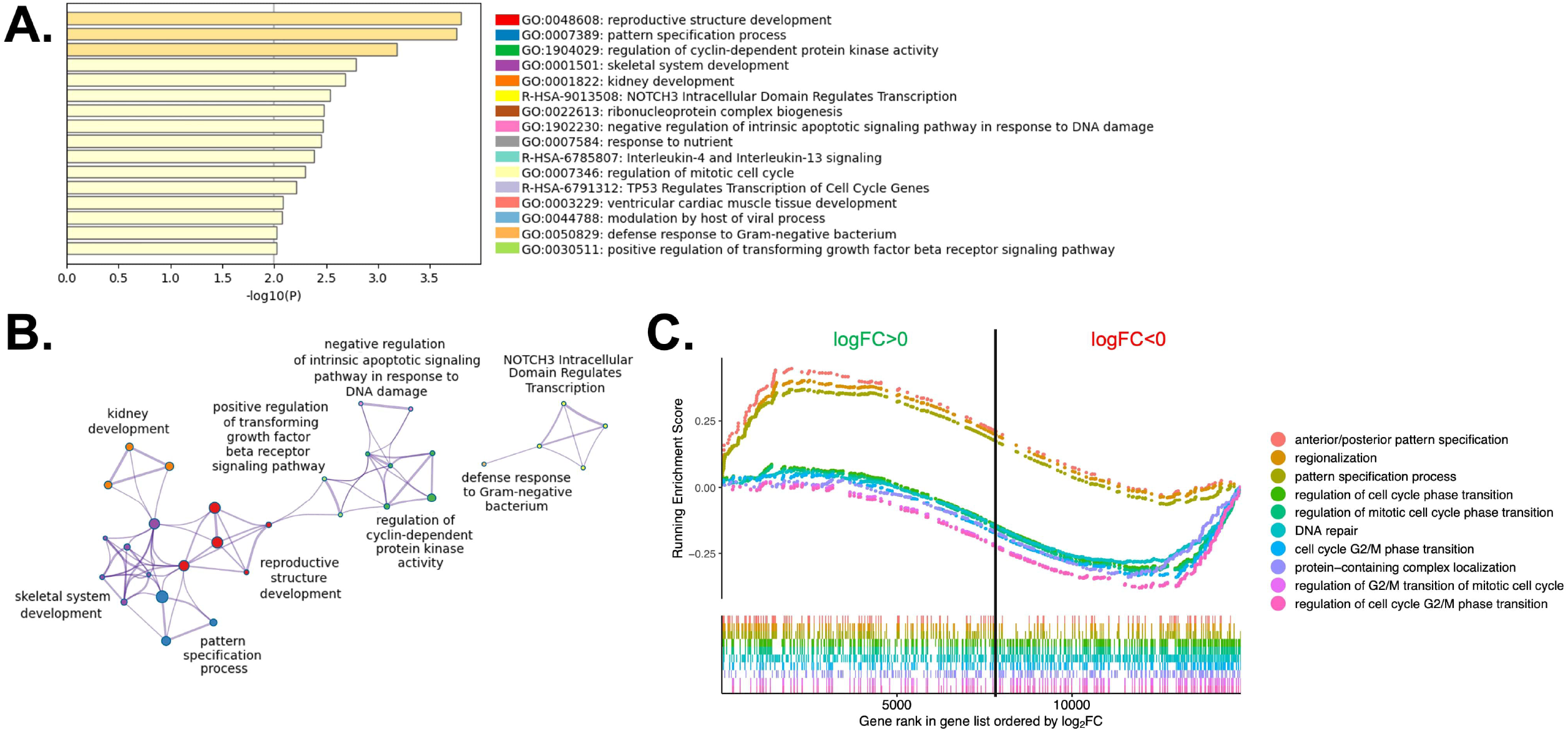
Gene Ontology analyses of Ferticult-to-Global DEGs at day-2. A. Overview of most significant GO terms from clusters of significant pathways over-represented in day-2 DEGs. Each term was selected by Metascape using a heuristic algorithm that selected the most informative term from clusters of proximal significant GO terms. B. Network of all GO significantly enriched terms, colored by their representative GO terms. Each node is an individual GO term and its size indicates the number of genes included in the GO term. Thick links connect GO terms of high similarity. GO terms with no intersection with other terms are not represented. C. Gene Set Enrichment Analysis (GSEA) of global gene expression changes in Ferticult-to-Global comparison. Only the top-10 biological processes are shown. Curves represent the running enrichment score of each GO term that reflects how each GO term contains over-(positive score) or under-expressed genes (negative score) with Ferticult. The running score enrichment increases by going through the overall gene ranking when a gene member of a GO term tested is met and decreases when a gene does not belong to the GO term tested. Bars at the bottom of the plot correspond to the rank at which the genes included in each GO term was assigned according to its log fold change with Ferticult. For example, concerning the genes involved in the anterior/posterior pattern specification pathway we observe highest density of genes with the largest positive logFC (pink curve). The vertical line separates genes with positive logFC (left side) from those with negative logFC (right side).

Expression changes may reflect chromatin modifications across the genome. Focusing on major genes involved in chromatin-based regulation such as DNA methylation, heterochromatin modulators, histone modifiers and remodeling complexes, we found two histone modifiers to be significantly down-regulated among the Ferticult-to-Global DEGs: the Aurora Kinase A gene *AURKA* (*p*=0.079, log2FC=-1.84), which regulates many aspects of mitosis, and *SETDB1* (*p*=0.073, log2FC=-2.92), which catalyzes trimethylation of the lysine 9 of histone H3 (H3K9me3) (Figure 4A). Additionally, focusing on imprinted genes, we only found the cyclin-dependent kinase inhibitor 1 (*CDKN1C)* gene among the 266 Ferticult-to-Global DEGs (Figure 4B). Finally, we also analyzed transposable elements and found three families to be differentially expressed between Ferticult and Global media (CR1-12_1Mi, LTR6A, LTR7C). Although the top-20 expressed transposable element families were not differentially expressed in any comparison, their expression was higher in Ferticult, which resulted in fold change intensities higher in Ferticult comparisons (Supplementary Figure S5A and S5B).

**Figure 4.**
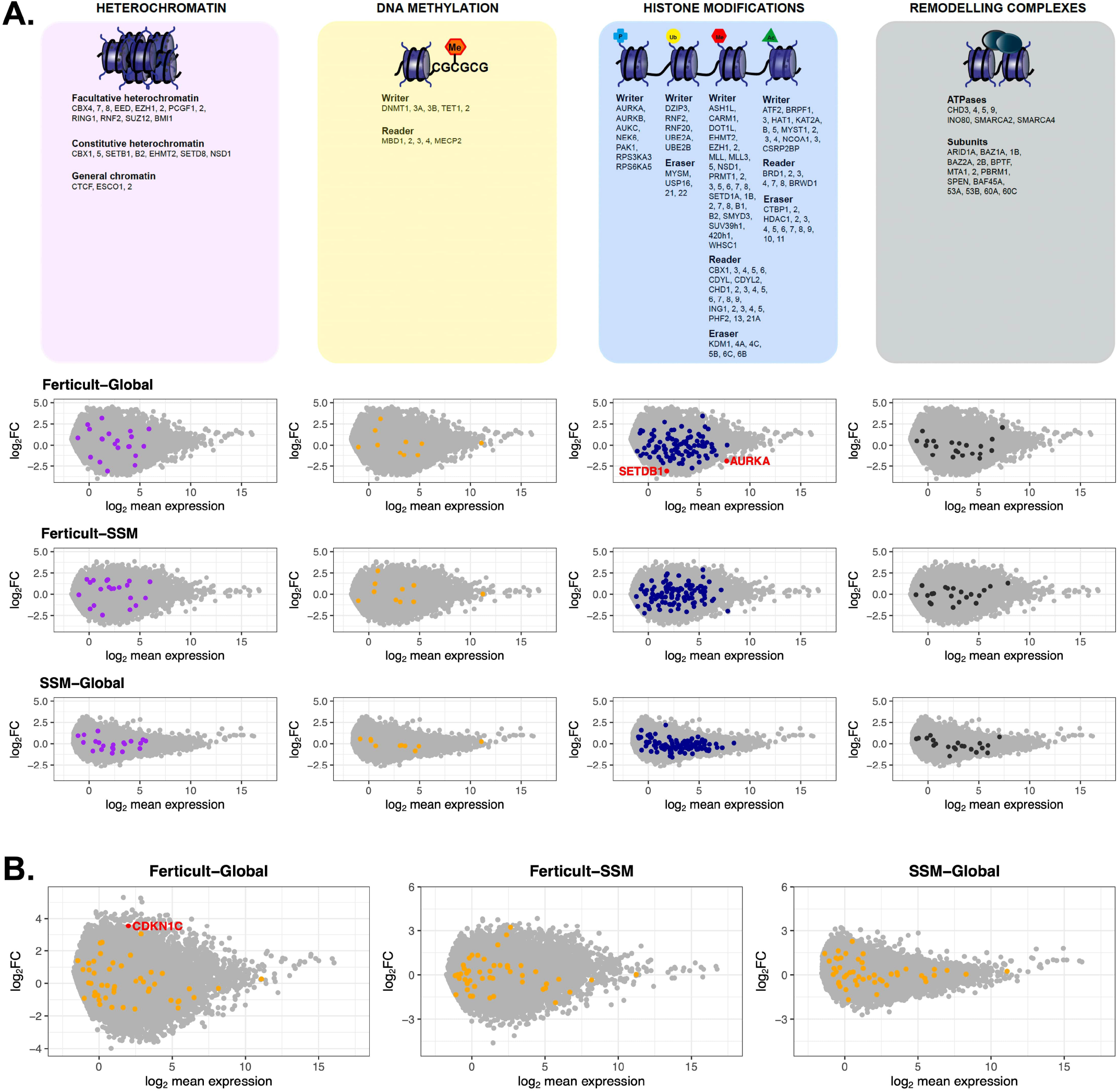
Expression differences of genes involved in chromatin-based mechanisms, imprinted genes and transposable elements between Ferticult, SSM and Global samples (day-2 embryos). A. List of major genes identified as involved in chromatin-based processes with their differential expression analysis. Log2 mean expression was calculated by taking the average log2(cpm+1) expression in compared culture media groups. B. Differential expression analysis focused on imprinted genes. Log2 mean expression was calculated by taking the average log2(cpm+1) expression in compared culture media groups. Mis-regulated imprinted genes (red dots), unchanged imprinted genes (orange dots), other expressed genes (gray dots).

### Monitoring 2 days culture medium-induced differences at the blastocyst stage

Given the transcriptomic differences of day-2 embryos resulting from amino-acid-free Ferticult medium over Global medium, we wanted to further analyze whether amino acid deprivation during the first two days of development may have extended effects on the transcriptome of blastocyst embryos. For that purpose, a second batch of embryos cultured until day-2 in Global (*n*=13) or Ferticult (*n*=7) media were selected for their strict identical embryo morphology, and subsequently cultured until the blastocyst stage, all in Global medium (Figure 1A). Performing differential expression analysis after single-embryo RNA-seq, we found 18 DEGs in blastocysts that were previously cultured in Ferticult medium until day-2 versus blastocysts cultured all along in Global : *ABCC6, AC008940*.*1, ACTL8, GPR143, H1FOO, HDC, HIST1H1A, KPNA7, NLRP4, NLRP13, PADI6, TUBB7P, TUBB8, TUBB8P7, TUBB8P8, TUBB8P12, WEE2* and *XAB2* (Figure 5A, 5B and Supplementary Table S4). These genes were all up-regulated with the Ferticult medium condition until day-2, with half showing a log2FC > 2.5, and the *XAB2* gene showing the highest overexpression score (>5.5) (Figure 5A). GO analysis indicated a functional link with meiotic cycle (Figure 5C). Importantly, none of the previous expression differences observed at day-2 remained significant at day-5. Circular plots showed that fold changes of gene differences observed at day-2 were largely minimized by day-5 (Figure 5D and 5E).

**Figure 5.**
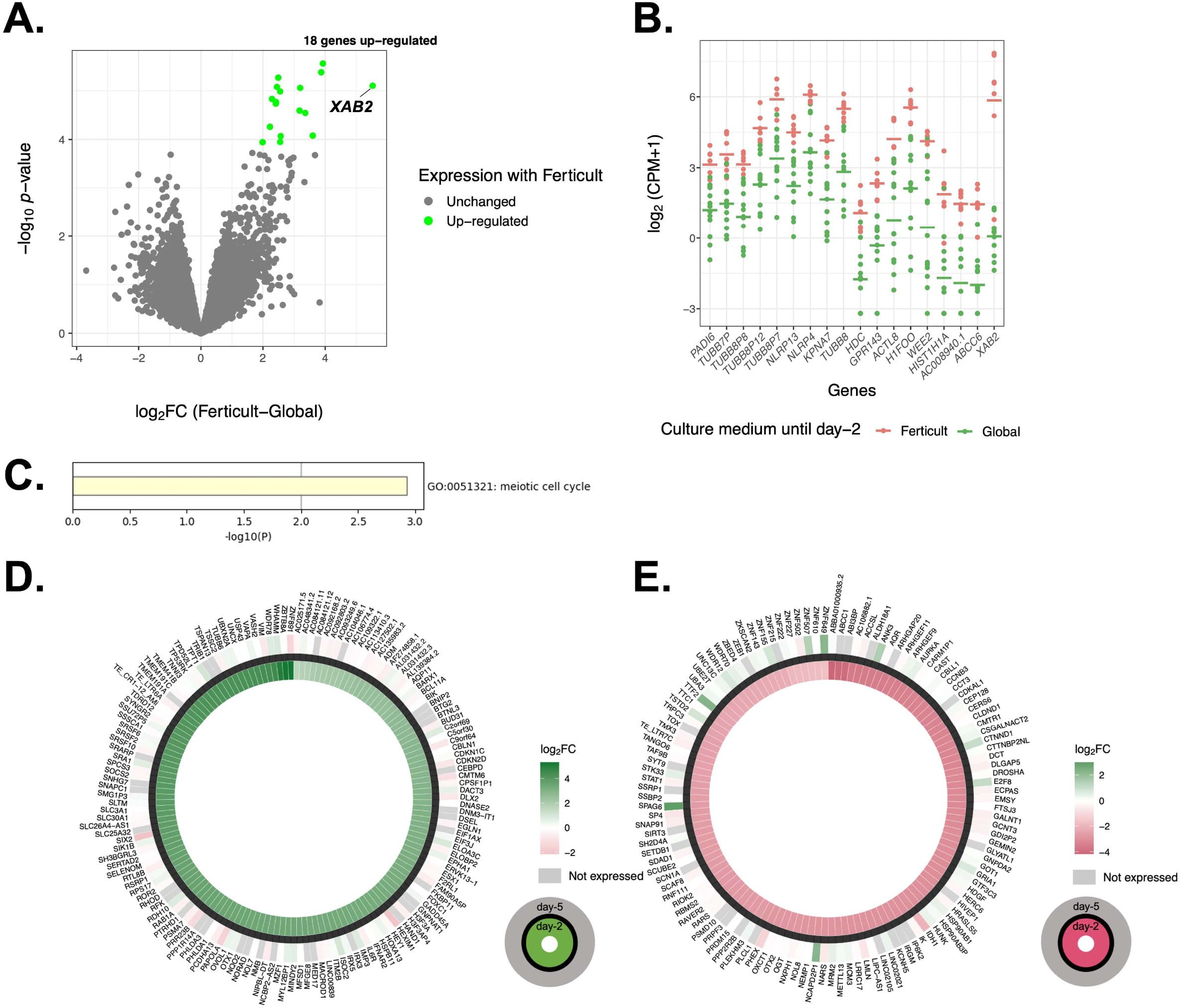
Differential expression analysis at day-5 of culture. A. Volcanoplot of differential gene expression between Ferticult and Global media. B. Dot plot of the expression of all 18 Ferticult-to-Global DEGs, ordered by ascending log2FC (from left to right). Group mean is represented by the line. Dots represent individual embryos. C. Significant pathways over-represented in day-5 DEGs. D. Circle plot of the expression of up-regulated Ferticult-to-Global DEGs at day-2 and their expression at day-5. Plot displays the log2 fold change of the 147 DEGs up-regulated with Ferticult at day-2 (interior layer) and the log2 fold change with Ferticult for the same genes at day-5 (exterior layer). Cells colored in grey correspond to genes that are not expressed at day-5. E. Circle plot of the expression of down-regulated Ferticult-to-Global DEGs at day-2 Global and their expression at day-5. Plot displays the log2 fold change of the 122 DEGs down-regulated with Ferticult at day-2 (interior layer) and the log2 fold change with Ferticult for the same genes at day-5 (exterior layer). Cells colored in grey correspond to genes that are not expressed at day-5.

### Effect of methionine supplementation

Methionine is an essential amino acid present in embryo culture media that serves as a precursor for protein synthesis and DNA methylation. We tested whether the addition of methionine at day-2 would impact the transcriptome of day-5 blastocysts cultured in Global by comparing 9 samples cultured in Global and 4 samples cultured in Global supplemented with methionine after day-2, from sibling embryos (i.e. a pair of embryos of each condition were coming from same couples) (Supplementary Figure S6A). Differential expression analysis revealed no significant DEG. However, GSEA was performed to assess whether the small insignificant differences might be related to any pathway. Thirty-two significant ontologies were identified upon methionine supplementation, which were mainly related to mitochondrial activity, oxidative phosphorylation and metabolism (Supplementary Figure S6B and Supplementary Table S5).

### Effects of culture medium on the transcriptional trajectory of early embryos

Early embryos undergo profound transcriptional changes during the first stages of development. In an attempt to address whether embryo culture medium may affect these sequential modifications of gene expression, we used a public dataset of 1,529 single-cell RNA-seq of human embryos from day-3 to day-7 (cultured in either CCM (Vitrolife) or G-1 Plus (Vitrolife) media) ^49^, which previously allowed delineating the transcription signature of each embryonic lineage and their dynamics during embryonic lineage segregation ^50,51^. Our objective was to identify genes whose expression dynamically changes across early embryonic development, in a continuous manner, independently of the embryo stage. For that purpose, we applied PHATE dimensionality reduction, a recently developed method that has been previously applied to human embryonic stem cells ^52^. We chose PHATE because of its ability to capture heterogeneity and reduce noise better than other dimensionality reduction methods ^52^. As the information geometry relies on diffusion dynamics, PHATE is especially suitable for early development ^52^. We then inferred existing lineages and pseudotime—a metric that could be interpreted as a timing distance between one cell and its precursor cell—using *slingshot*, a method adapted for branching lineage structures in low-dimensional data. On the public scRNA-seq dataset, we were able to identify 8 clusters by using k-means clustering to group of cells with high transcriptomic similarities. We also identified three distinct lineages (Supplementary Figure S7A) that shared the same structure when considering cells from clusters 1 to 5, but separated into clusters 6, 7 and 8. Using the cell classification adopted in previous studies ^49,51^, the three lineages corroborated with the demarcation into epiblast (EPI), primitive endoderm (PrE) and trophectoderm (TE) cells (Supplementary Figure S7B and S7C). Each cell was then assigned a pseudotime to reflect its “transcriptomic age” along each of the three lineages (Supplementary Figure S7D). Along this inferred pseudotime, we identified 1,110 genes with a dynamic expression pattern (Figure 6A).

**Figure 6.**
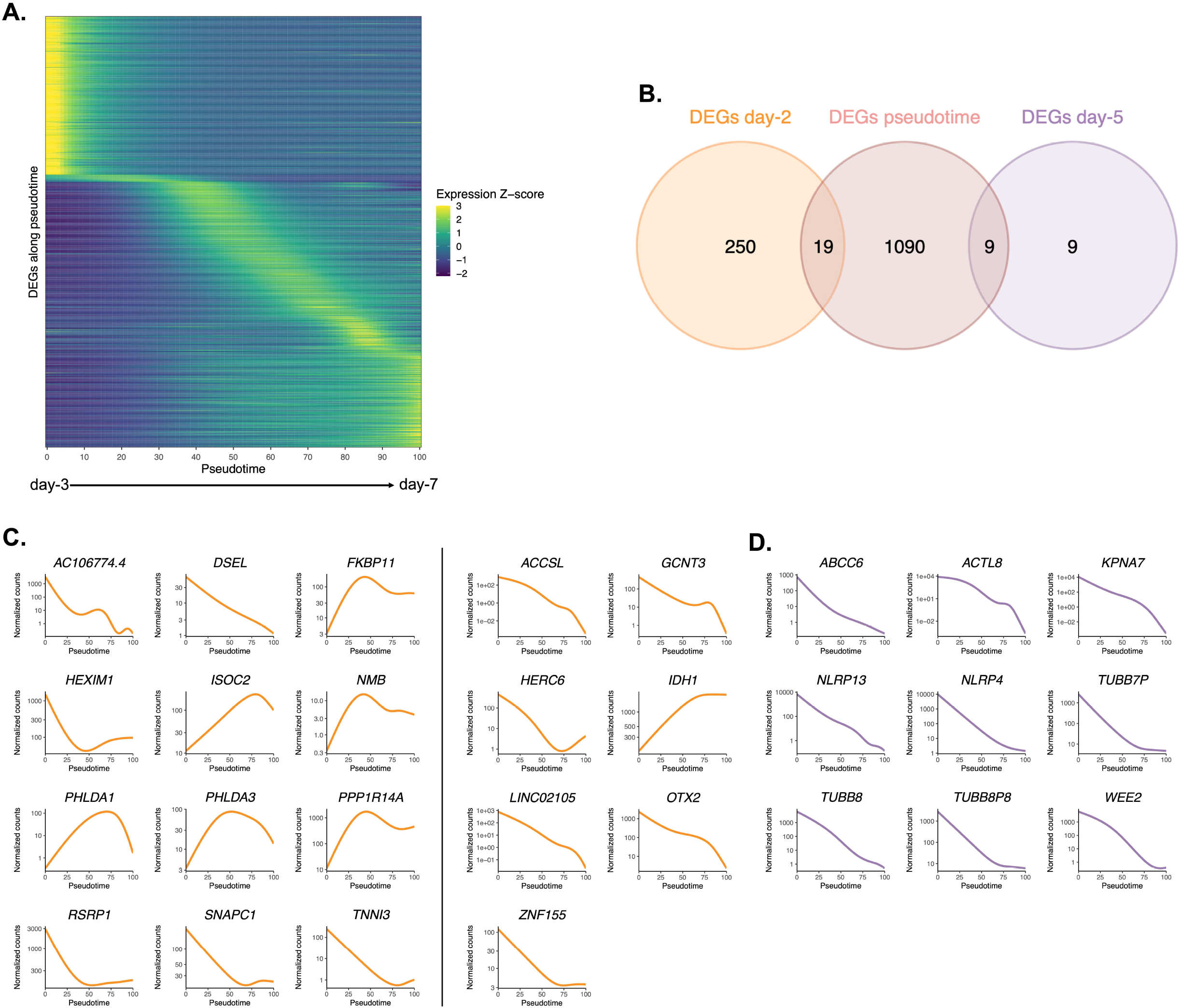
Pseudotime differential expression during human preimplantation development. A. Heatmap of expression of the 1,110 genes that were found differentially expressed along pseudotime (from public datasets established from 8-cell stage ^49^), ordered by timing of peak expression (arbitrary unit). Expression Z-score : Z-score of TMM-adjusted cpm. B. Venn diagram of the number of DEGs along pseudotime that are differentially expressed between Ferticult and Global at day-2 and day-5 of embryonic culture. C. Dynamic of expression of the 19 day-2 Ferticult-to-Global DEGs that are differentially expressed along pseudotime. The curve corresponds to the NB-GAM fitted normalized counts (TMM-adjusted cpm). Left panel corresponds to genes up-regulated with Ferticult. Right panel corresponds to genes down-regulated with Ferticult. D. Expression dynamics of the 9 day-5 Ferticult-to-Global DEGs that are differentially expressed along pseudotime. The curve corresponds to the NB-GAM fitted normalized counts (TMM-adjusted cpm).

Considering the question of the impact of culture medium, we crossed these 1,110 dynamic genes with our list of DEGs identified at day-2 and day-5 in the Ferticult-to-Global comparison. Remarkably, 19 out of 269 DEGs at day-2 (transposable elements included) and 9 out of 18 DEGs at day-5 showed dynamic expression changes across pre-implantation pseudotime, meaning that the choice of culture medium has an impact on the expression of genes that are dynamically regulated during early development (Figure 6B). Temporal expression of those genes is shown in Figure 6C and 6D. The 9 DEGs at day-5 that are also differentially expressed along pseudotime displayed declining expression over embryo development in the reference pseudotime (Figure 6D), which was confirmed using an independent public scRNA-seq dataset obtained from oocyte to late blastocyst^47^ (Figure 7). When considering our own datasets, embryos continuously cultured in Global up to day-5 showed also this declining trend of expression from day-2 to day-5, with levels that were congruent with the reference dataset (Figures 7). However, day-5 embryos previously cultured in Ferticult until the 4-cell stage showed over-expression, suggesting that these embryos retained abnormally high levels for their embryonic stage. On average, these blastocyst embryos showed expression levels that were closer to morula stages.

**Figure 7.**
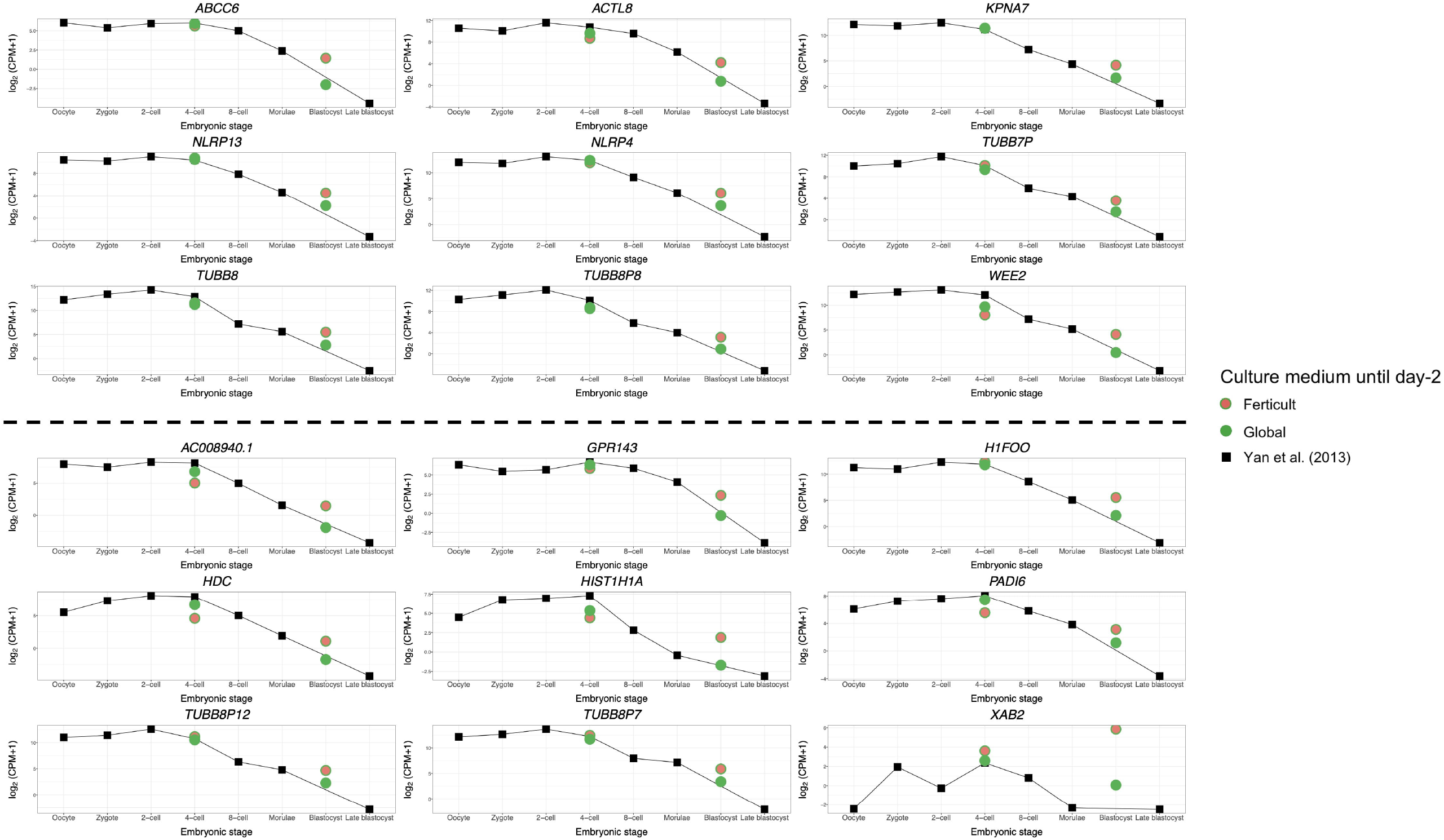
Expression dynamics of day-5 Ferticult-to-Global DEGs from oocyte to blastocyst stage. Public scRNA-seq datasets ^47^ obtained from oocyte to blastocyst stage were plotted as reference level of expression of early embryonic genes. Each colored point represents the mean log2(cpm+1) value of all samples for each culture medium group. Day-2 embryos were represented at the 4-cell stage. Day-5 embryos were represented at the blastocyst stage. Genes above the dashed line were found differentially expressed along pseudotime.

## Discussion

We provide here an in-depth characterization of the transcriptomic effects exerted by different IVF media on human embryos after 2 and 5 days of culture. It yields several insights into how culture medium composition can induce transcriptomic responses as an adaptation of the embryo to its micro-environment, pre- and post-compaction. In line with the importance of our research question, two of the media we tested are no longer used due to their underperformance in IVF outcomes (preimplantation embryo development and pregnancy rates).

First, we focused on embryos cultured until day-2, as this pre-compaction period is likely to be sensitive to environmental stressors. The most pronounced transcriptional divergence was between Ferticult and Global, with 266 DEGs. Nineteen of these DEGs could have a key role in early development, based on their dynamic expression changes across development but also on their association with GO terms related to essential developmental pathways, and were mostly up-regulated in Ferticult. Our results are congruent with two previous microarray studies that measured transcriptomic differences between embryos cultured in two culture media (G5 and HTF) ^53^.

Imprinted genes are candidates for high susceptibility to environmental conditions and disruption of imprinted expression has been linked to developmental pathologies in Human 54. Accordingly, animal studies indicated that some embryo culture media were associated with hypomethylation of maternally expressed genes (such as *H19* and *SNRPN*) resulting in aberrant biallelic expression ^38,55^. In our study, only one imprinted gene, *CDKN1C*, was up-regulated after 2 days in culture in Ferticult compared to Global. *CDKN1C* is a key regulator of cell growth and proliferation, and aberrant expression is observed in syndromes with overgrowth, tumor predisposition and congenital malformations, such as Beckwith-Wiedemann syndrome ^56^. We also found the expression of *SETDB1*, which is involved in histone methylation, was down-regulated in Ferticult samples. These elements may reflect direct and indirect influences of the culture medium on the embryonic epigenome.

The link between culture medium composition and transcriptomic effects is still unclear, but transcriptional changes may reflect an adaptation to a sub-optimal environment. For those reasons, we further tested whether the differences observed at day-2 in Ferticult over Global were maintained later on, at the blastocyst stage, after being cultured in Global, which is considered more suitable because of its amino acid enrichment ^36^. Only 18 DEGs were retrieved, and importantly, none of the early differences observed at day-2 were conserved at this later stage. Notably, the misregulation of *AURKA, SETDB1* and the imprinted *CDKN1C* gene observed at day-2 no longer existed at day-5. The original transcriptional changes may have not persisted because, in post-compaction, the embryo acquires increasing ability to respond to different environments and to correct transcriptional errors ^57^. A second hypothesis is that the Global medium composition itself may have allowed the embryo to recover a favorable transcriptomic landscape. Finally, we cannot rule out that only viable embryos were able to develop until the blastocyst stage and only embryos with functional abilities were therefore selected in our analysis.

Nevertheless, our analysis of genes that are differentially expressed along pseudotime brought evidence that the use of distinct culture media prior to compaction can alter the sequential gene expression changes linked to later embryo development. Genes activated or down-regulated at the wrong time may impact the development and cause lasting effects ^58,59^. Notably, Ferticult was associated with systematic over-expression of genes at day 5. It is therefore possible that two days of culture in Ferticult induce a developmental delay, which could be manifested by a delay in maternal RNA clearance, for example. Accordingly, two of the 18 DEGs at day-5 are maternal effect genes (*PADI6* and *TUBB8*)^60^, which may indeed reflect longer retention of maternal transcripts due to developmental delay. *PADI6* is a member of the subcortical oocyte complex ^61,62^, while *TUBB8* is the major constituent of the oocyte meiotic spindle assembly in primates ^63^.

Nevertheless, our pseudotime analysis of genes differentially expressed over the course of development also identified genes that are thought to be involved in the transition from early to later embryonic stages, such as *ABCC6, ACTL8, KPNA7, NLRP4, NLRP13, TUBB7P, TUBB8P8*, and *WEE2. TUBB7P* and *TUBB8P8* encode beta-tubulins of major importance in cell division and morphology. *KPNA7* (karyopherin subunit alpha 7) is involved in nuclear protein transport ^64^ and *Kpna7*-deficient mice fail to develop to blastocyst stage or show developmental delays ^65^. Whether *ABCC6, ACTL8, NLRP4, NLRP13* and *WEE2* are involved in early embryogenesis remains unknown. Additionally, *XAB2* (XPA binding protein 2), whose expression does not appear to be stage-specific, was particularly high in Ferticult (log2FC>5.5). XAB2 plays a role in DNA repair ^66^ and is required for embryo viability ^67,68^. Activation of DNA repair mechanisms may be reflective of stress conditions experienced by pre-implantation embryos. Because embryos were cultured in the same medium after day-2 in our study and because the embryo is transcriptionally silent until EGA, we can hypothesize that the blastocyst transcriptome was influenced by alterations that occurred pre-compaction.

Finally, we investigated whether adding methionine in the culture medium, an essential amino acid whose concentration varies greatly between commercial media ^69^, could affect embryo gene expression. Methionine is a precursor of S-adenosylmethionine, a key component in the one-carbon metabolism and methylation processes ^70^. Methionine is necessary for proper embryo development but in excessive concentration, it could negatively affect embryo abilities, as demonstrated in several animal models ^71–73^. Reassuringly, we did not identify any DEGs in sibling embryos cultured in Global until day-5, with or without methionine supplementation, suggesting that excessive methionine concentration from day-2 did not have major influence on the blastocyst’s transcriptome. Despite the lack of significant differences, GSEA revealed activation of several mitochondrial activities and metabolic processes, in agreement with the stimulatory effect of methionine on mitochondrial respiration ^74^. Analyzing the early effects of methionine addition before EGA could be important.

Evidence that embryo culture medium can impact gene expression has long been described in animals ^36,39,40^. Interestingly, in pig, adding -reproductive fluids during *in vitro* culture allows producing blastocysts with closer chromatin and transcriptomic profiles compared to natural conditions ^75^. It will be important to develop culture media closer to natural fluid even if we showed that the embryo is highly adaptable to different conditions. Additionally, if this study is reassuring, we might not forget that many other processes in the IVF laboratory environment constitute environmental stressors (temperature, pH, co-culture, light, oxygen tension, manipulation). In our design, culture conditions other than culture media were identical for all samples, but a gamete or an embryo that has been exposed to a stressful condition might be even more vulnerable to other stressors.

For the first time in Human, we employed single-embryo RNA-seq on day-2 and day-5 embryos to assess to what extent different culture conditions might affect the developing embryo at the molecular level. Even though marked transcriptomic differences were observed between culture media at day-2, when embryos totally deprived in amino acids during their first days of development were returned into favorable culture conditions, these differences were mitigated by the blastocyst stage. The few differences observed at day-5 may be attributed to a delay in molecular processes specific to the use of one medium. Altogether, this study emphasizes the abilities of the embryo to recover an appropriate transcriptome post-compaction. Consecutively, to rule out potential long-lasting epigenetic effects, it would be important to investigate whether the methylome also adapts to different media formulations.

## Materials and methods

### Ethics statement

This research was authorized by the National Biomedicine Agency (Legal decision published in the Journal Officiel under the reference JORF N°0233-the 6th October 2016 and extended under the reference JORF n°0303-the 31th December 2019).

### Embryo selection-experimental design

We used embryos donated for research by couples and cryopreserved at the Reproductive Lab of Dijon Hospital. All embryos used in the current study were cryopreserved individually 2 days post fertilization and had been *in vitro* cultured in different culture media (Global medium, LifeGlobal; FertiCult IVF medium, FertiPro; SSM, Irvine Scientific). For the first question of the project, we analyzed only embryos cultured in these three different culture media but with identical morphological criteria i.e. at 4-cell stage with less than 15% of anucleate fragments and regular cleavage (Figure 1). For the second question, we selected day-2 cryopreserved embryos with identical morphological criteria as described above from embryo cohorts cultured up to day-2 either in FertiCult or Global, which were afterwards cultured up to day-5 in Global. Finally, for methionine supplementation, we included day-2 cryopreserved embryos from embryo cohorts from the same patients with at least four embryos with identical morphological criteria. After thawing, embryos were randomly cultured in Global medium without or with methionine supplementation (200µM; concentration nearly thrice that in Global medium).

Embryos were thawed according to the strict protocol routinely used in human IVF-clinic to maintain as much as possible their integrity ^76^. Immediately after thawing, embryos were transferred into preequilibrated Embryoslides (Unisense Fertilitech, Vitrolife) with 25 μl of culture medium and covered with 1.2 mL of oil (Nidoil, Nidacon). They were cultured up to the blastocyst stage at 37°C and a tri-gaz atmosphere (6%CO2, 5%O2, 89% N2). According to the classification of Gardner and Schoolcraft (1999) ^77^, only blastocysts with at least B2 blastocoel cavity without lysis were analyzed. We also paid attention to use the same batches of culture media in all experiments.

### Single-embryo RNA sequencing

A previously described scRNA-seq method was applied to single embryos ^46^. Briefly, zona pellucida-free embryos (after using acidic Tyrode’s solution) were individually placed in a lysis buffer containing 1.35mM MgCl_2_ (4379878, Applied Biosystems), 4.5 mM DTT, 0.45% Nonidet P-40 (11332473001, Roche), 0.18U/mL SUPERase-In (AM2694, Ambion) and 0.36U/mL RNase-inhibitor (AM2682, Ambion). Then, we performed a reverse transcription reaction (SuperScript III reverse transcriptase - 18080-044, Invitrogen, final concentration : 13.2 U/mL) and poly(A) tailing to the 3’ end of the first-strand cDNA (by using terminal deoxynucleotidyl transferase - 10533-073, Invitrogen, final concentration : 0.75U/mL). After the second-strand cDNA synthesis, 20 and 18 cycles (at day-2 and day-5, respectively) of PCR were performed to amplify the embryo cDNA using the TaKaRa ExTaq HS (TAKRR006B, Takara, final concentration : 0.05U/mL) and IS PCR primer (IDT, final concentration : 1mM). Following purification with Zymoclean Gel DNA Recovery Kit (ZD4008, Takara), product size distribution and quantity were assessed on a Bioanalyzer using an Agilent 2100 high-sensitivity DNA assay kit (5067-4626, Agilent Technologies). The library preparation (KAPA Hyper Plus Library prep kit) and sequencing was performed by the ICGex - NGS platform (Institut Curie) on HiSeq 2500 for day-2 embryos and on NovaSeq 6000 Illumina sequencer for day-5 embryos for 100-bp paired-end sequencing (Supplementary Table S1).

### Data pre-processing and quality control

We computed sequencing quality checks with FastQC v0.11.9 and trimming of adapters and low-quality sequences using TrimGalore! v0.6.6. Paired-end reads alignment was performed onto Human reference genome (hg38) with STAR v2.7.9a ^78^ reporting randomly one position, allowing 6% of mismatches. Following previous recommendations ^79^, repeat annotation was downloaded from RepeatMasker and joined with basic gene annotation from Gencode v19. The merged file was used as input for quantification with featureCounts v2.0.1. Genes with a minimum of count per million (cpm) > 1 in at least 4 samples were retained for further analysis. Principal component analyses were implemented with PCAtools v2.8.0 on log2(cpm+1) for all genes for single datasets and common genes for multiple datasets, except the 10% genes with lowest variance.

### Differential expression analysis

Differential expression analysis was performed using *edgeR*’s normalization (v3.38.1) combined with *voom* transformation from *limma* package v3.52.1. *P*-values were computed using *limma* and adjusted with the Benjamini-Hochberg correction. Genes were declared as differentially expressed if FDR<0.1.

### Gene Ontology and Gene Set Enrichment Analysis

We used Metascape v3.5 to calculate and visualize over-representation of gene ontologies in our list of differentially expressed genes ^80^. Metascape applies hypergeometric test and FDR correction to identify ontology terms that comprise significantly more genes in a given gene list that what would be expected with a random gene list. For each gene list tested, we provided an appropriate background gene list corresponding to all expressed genes in all samples for a given experiment. We selected “Express Analysis” to capture relevant gene annotation from multiple sources (GO, KEGG, Reactome, canonical pathways, CORUM). The P-value cutoff was kept at 0.01.

Gene Set Enrichment Analysis was implemented with the *clusterProfiler* R package (v4.4.1) setting adjusted p-value significance threshold at 0.05. Beforehand, imputed gene lists were pre-ranked by logFC.

### Processing of public single-cell RNA-seq datasets in human embryos

We compared our data with three early embryos single-cell RNA-seq studies ^47–49^. Reads alignment and quantification was executed as described previously onto raw reads downloaded from European Nucleotide Archive (study accession PRJNA153427, PRJNA189204 and PRJEB11202).

### Trajectory inference and pseudotime computing

After pre-processing, read counts data from all 1529 cells from Petropoulos *et al*. was pre-clustered and normalized with *scran* and *scater* packages (v1.24.0) after removing lowly expressed genes. Next, *Seurat* (v4.1.1) was used to scale the data and regress variable on total RNA. PHATE dimension reduction was applied with *phateR* v1.0.7 embedding three dimensions and used as input for pseudotime computing and trajectory inference with *slingshot* v2.4.0.

### Differential expression along pseudotime

We used *tradeSeq* v1.10.0 ^81^ to fit negative binomial generalized additive model (NB-GAM) for each gene. After examining diagnostic plots of the optimal number of knots (k) according to Akaike Information Criterion (AIC), k was set to 6 as optional parameter in the NB-GAM model. We selected differentially expressed genes along pseudotime with *associationTest()* function if *p*-value<0.05 and meanLogFC>2. This function relies on Wald tests to assess the null hypothesis that the expression of a gene is constant along pseudotime.

### Code and data availability

All data generated in this study have been deposited in the Gene Expression Omnibus (GEO) under accession number GSE212811 (https://www.ncbi.nlm.nih.gov/geo/query/acc.cgi?acc=GSE212811). All code processed in this study can be accessed on https://github.com/BastiDucreux.

## Acknowledgements

We thank Fuchou Tang for hosting Patricia Fauque in his lab to learn the scRNA-seq method, Nicolas Lieury for his assistance in preparing and obtaining the samples and Maud Carpentier of the “Direction de la Recherche Clinique et de l’Innovation” of Dijon University Hospital for the financial management of the study. We acknowledge the ICGex NGS platform of Institut Curie (supported by grants ANR-10-EQPX-03, Equipex and ANR-10-INBS-09-08, France Génomique).

## Funding

This work was supported by funding from the “Agence Nationale pour la Recherche (“CARE”-ANR JCJC 2017)”.

## Author contributions

P.F., J.B. and B.D. took primary responsibility for conceptualization and investigation. P.F. and R.P-P were responsible for the methodology. M.G., J.B. and P.F. were involved in resources, experiments and visualization. B.D. and A.T. did the data curation and formal analysis. B.D and P.F. were involved in original draft preparation. D.B. participated to review and editing.

## Competing interests

The authors declare no competing interests.

## Supplementary Figure legends

**<u>Supplementary Figure S1</u>. Overlap of expressed genes in our study with Yan et al. and Xue et al. data**.

A. Table and Venn diagram indicating the number of expressed genes that overlap between studies at day-2.

B. Venn diagram indicating the number of expressed genes that overlap between studies at day-5.

**<u>Supplementary Figure S2</u>. Principal component analysis of Yan et al**., **Xue et al. and this study’s datasets according to the normalized expression (log2(cpm+1)) of all expressed genes**.

**<u>Supplementary Figure S3</u>. Pairwise comparison of the levels of expression of all genes expressed between Ferticult, Global and SSM groups**. R: Spearman’s correlation coefficient.

**<u>Supplementary Figure S4</u>. Detailed analysis of significant GO terms significantly over-represented in DEGs between Ferticult and Global at day-2**.

A. Overview of most significant GO terms from clusters of significant pathways over-represented in day-2 differentially expressed genes (DEGs).

B. Network of all GO significantly enriched terms, colored by their representative GO terms. Three clusters of GO terms interaction were identified. Each node is an individual GO term and its size indicates the number of genes included in the GO term. Thick links connect GO terms of high similarity.

C. Detailed expression of DEGs between Ferticult and Global at day-2 involved in clusters of enriched pathways.

**<u>Supplementary Figure S5</u>. Differential expression analysis at day-2 focused on top-20 expressed transposable elements for all groups**.

A. Log2 fold change of transposable elements in all comparisons between culture media.

B. Mean expression of transposable elements in all culture media groups. Log2 mean expression was calculated by taking the average log2(cpm+1) expression.

**<u>Supplementary Figure S6</u>. Effects of a methionine supplementation in the culture medium at day-2**.

A. Study design of the experiment.

B. Gene Ontologies affected by methionine supplementation in embryo culture media (assessed via GSEA). Only the top-10 biological processes are shown.

**<u>Supplementary Figure S7</u>. PHATE reduction of Petropoulos’ (2016) embryo scRNA-seq data**. Each point represents one of the 1529 cells of the dataset. Curves indicate trajectory of the three inferred lineages by *slingshot*.

A. Colored by k-means clusters identified in this study. Cells were grouped into 8 clusters according to their global expression of expressed genes. This clustering was used as input in *slingshot* to depict the global structure of underlying embryonic lineages.

B. Colored by Meistermann et al. lineage inference. Authors applied UMAP dimensionality reduction on module eigengenes identified with WGCNA (Langfelder and Horvath, 2008) to cluster cells according to their association with gene expression signatures specific of developmental stages and lineages.

C. Colored by Petropoulos et al. original lineage inference. Authors used PCA dimensionality reduction and to infer embryonic lineages of cells after day-5.

D. Colored by pseudotime inferred with *slingshot* in this study (arbitrary unit). Pseudotime of the second lineage has been chosen for purpose of visualization. Earliest embryonic stages of the dataset are grey colored while dark red colored points are indicative of latest cells timepoints.

## Supplementary Table Legends

**<u>Supplementary Table S1</u>. Single-embryo RNA-sequencing statistics for all samples**.

**<u>Supplementary Table S2</u>. Differential expression analysis of all groups at day-2**. Genes highlighted in yellow reached significance (*p*<0.1).

**<u>Supplementary Table S3</u>. Gene Set Expression Analysis results for the Ferticult-Global comparison at day-2**.

**<u>Supplementary Table S4</u>. Differential expression analysis of all groups at day-5**.

**<u>Supplementary Table S5</u>. Gene Set Expression Analysis results for the methionine supplementation analysis**.

## Notes

### Competing Interest Statement

The authors have declared no competing interest.

