## Supplementary figures and images for "Assessing the influence of distinct IVF culture media on human pre-implantation development using single-embryo transcriptomics"

### Supplementary Figure S1

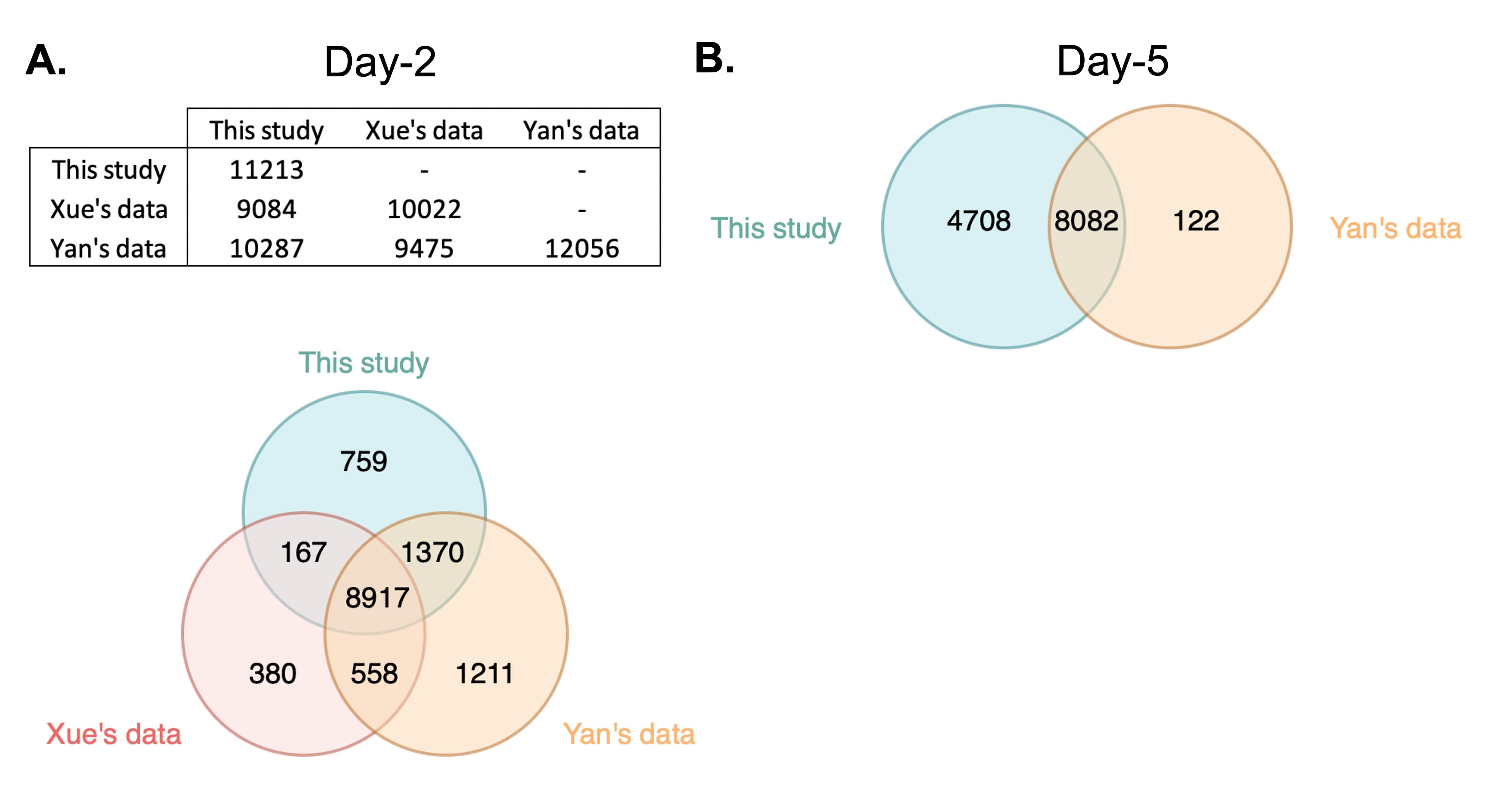

### Supplementary Figure S2

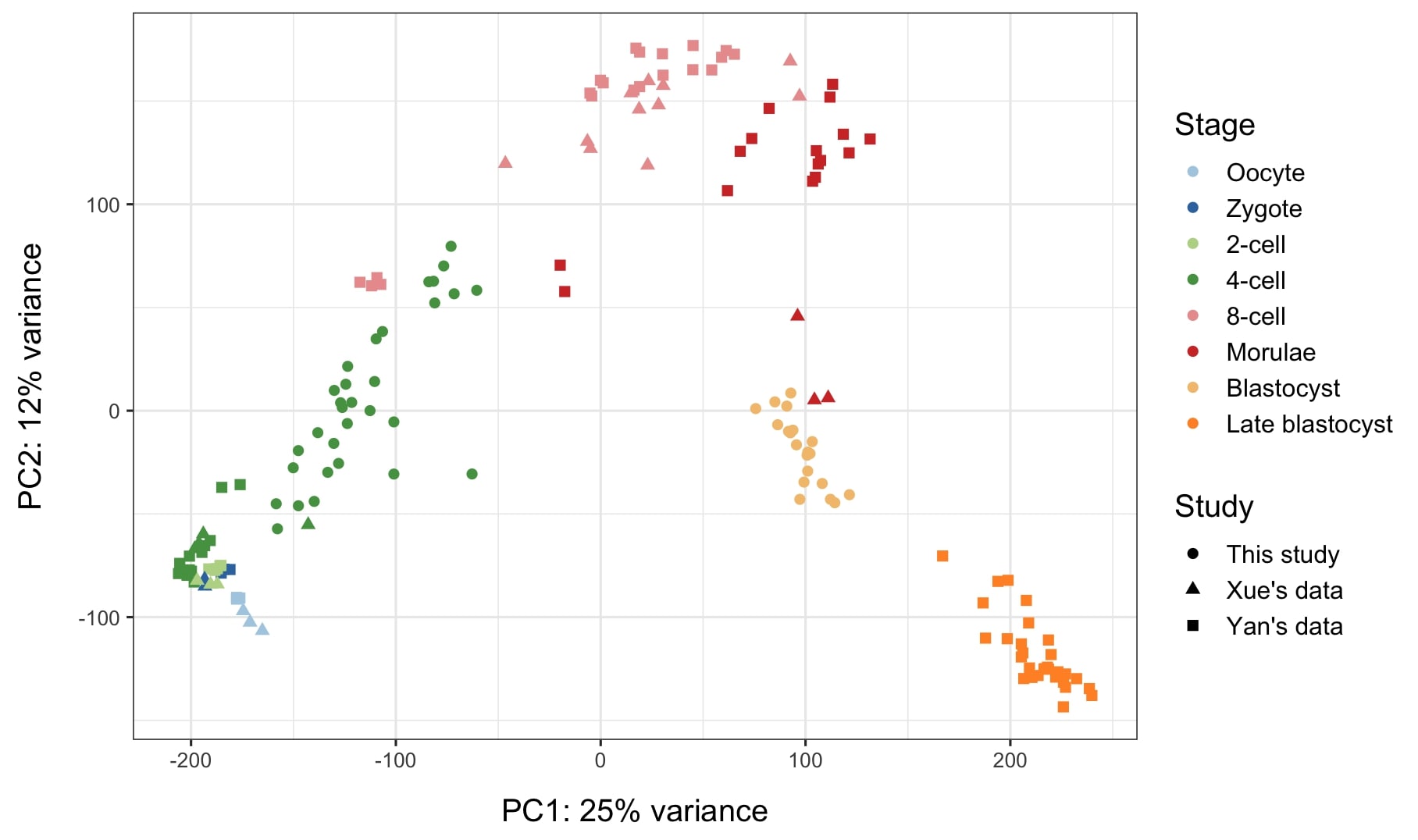

### Supplementary Figure S3

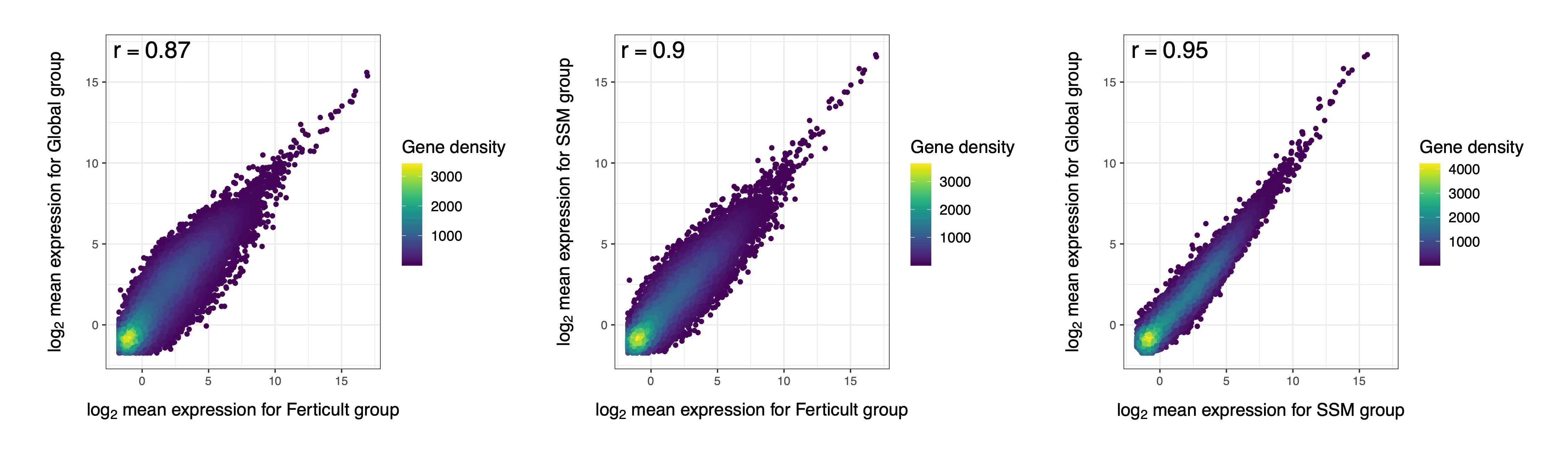

### Supplementary Figure S4

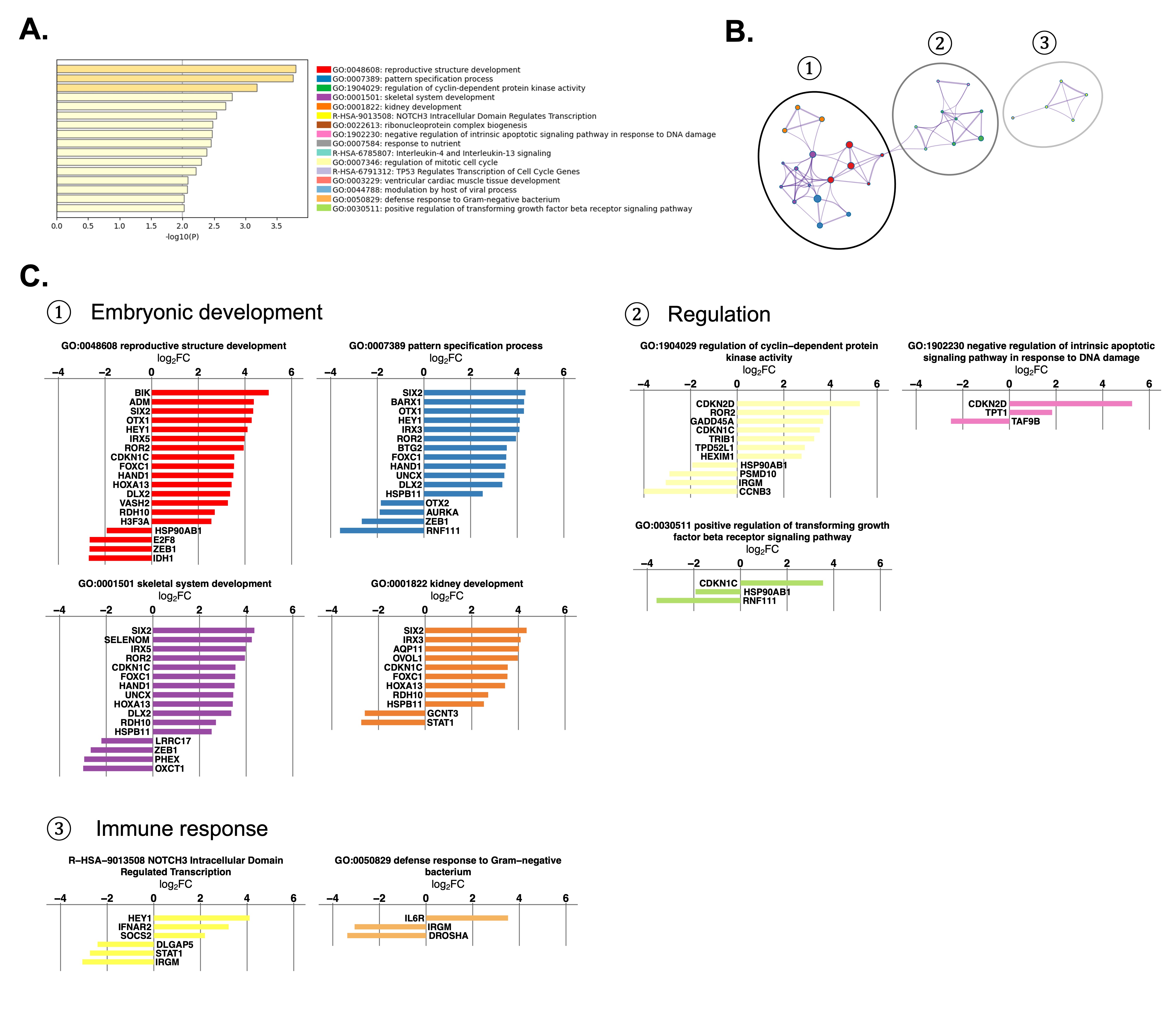

### Supplementary Figure S5

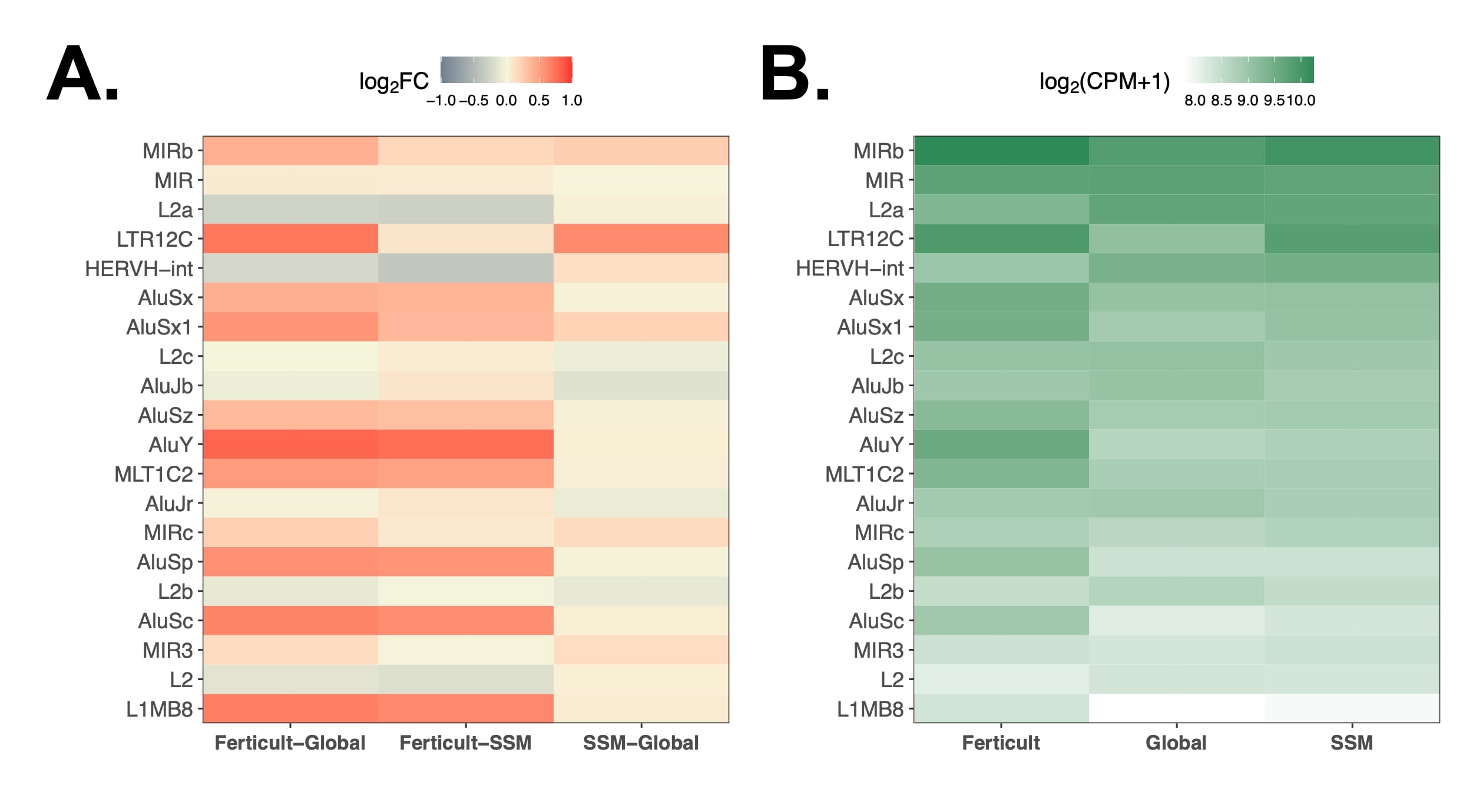

### Supplementary Figure S6

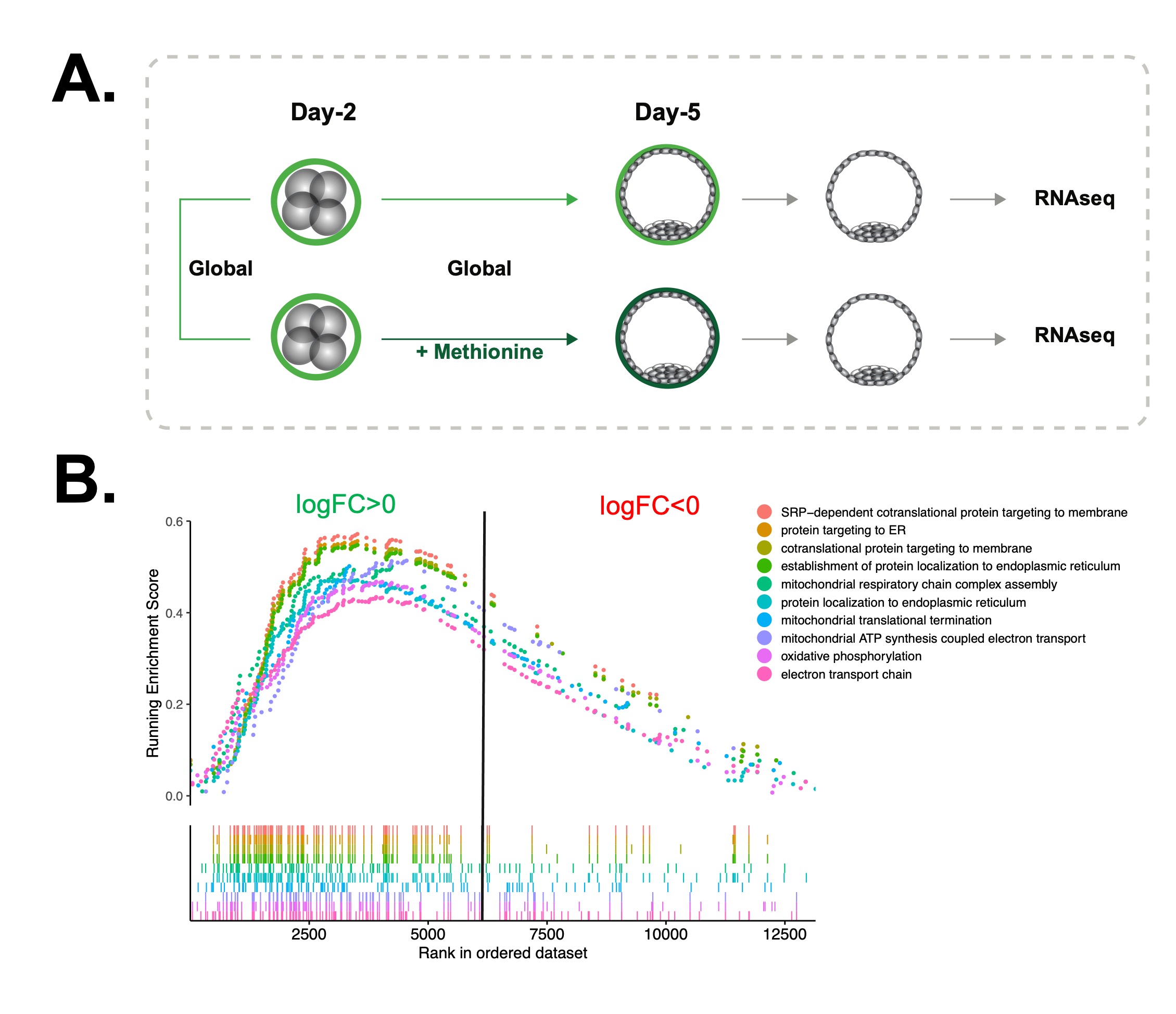

### Supplementary Figure S7

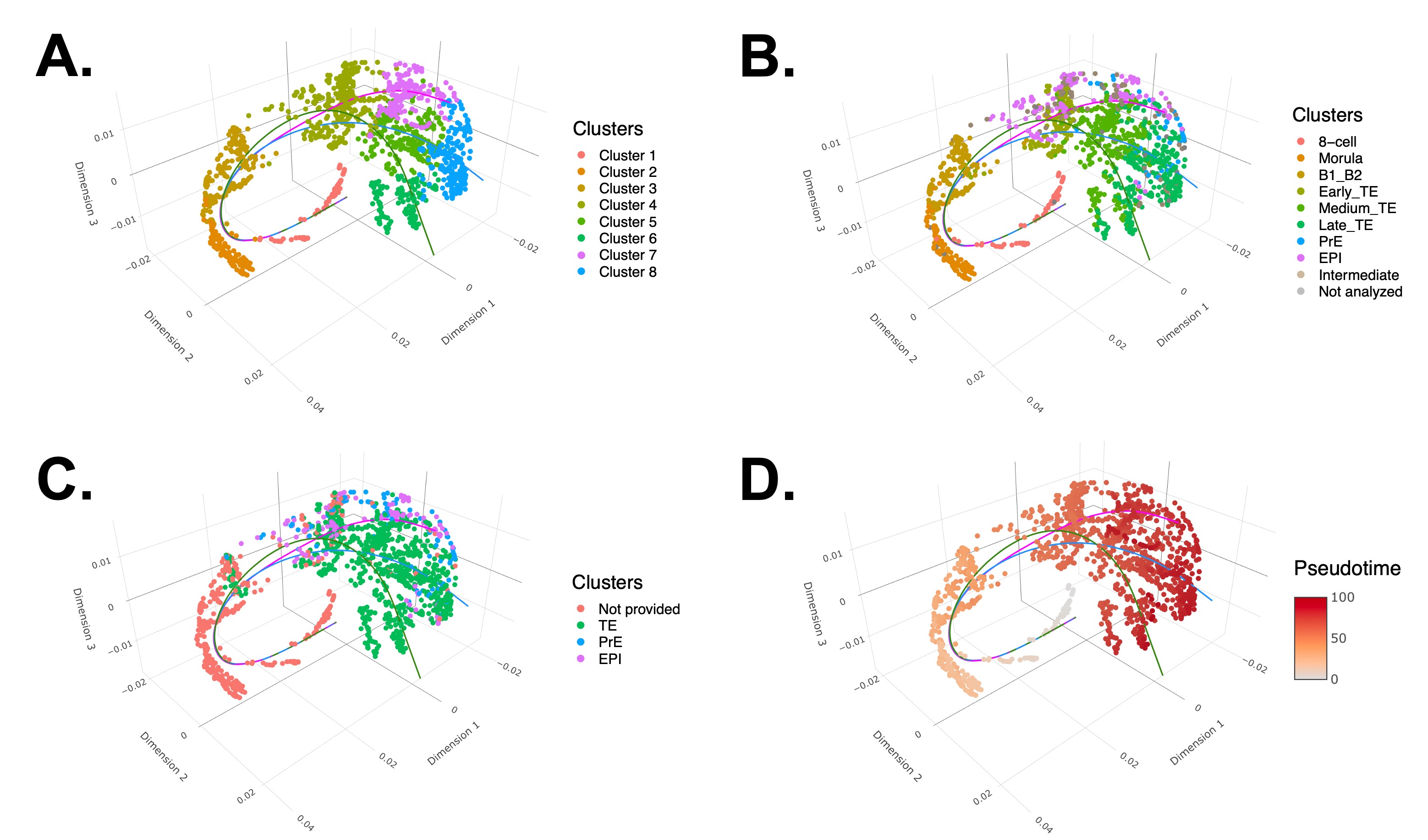
